# The *Drosophila melanogaster* pheromone Z4-11Al is encoded together with habitat olfactory cues and mediates species-specific communication

**DOI:** 10.1101/083071

**Authors:** Sebastien Lebreton, Felipe Borrero-Echeverry, Francisco Gonzalez, Marit Solum, Erika Wallin, Erik Hedenström, Bill S. Hansson, Anna-Lena Gustavsson, Marie Bengtsson, Göran Birgersson, William B. Walker, Hany Dweck, Paul G. Becher, Peter Witzgall

**Author notes:** Correspondence: Peter Witzgall, SLU, Box 102, 23053 Alnarp, Sweden.

## Abstract

Mate recognition in animals evolves during niche adaptation and involves habitat and social olfactory signals. *Drosophila melanogaster* is attracted to fermenting fruit for feeding and egg-laying. We show that, in addition, female flies release a pheromone (*Z*)-4-undecenal (*Z*4-11Al), that elicits flight attraction in both sexes. The biosynthetic precursor of *Z*4-11Al is the cuticular hydrocarbon (*Z*,*Z*)-7,11-heptacosadiene (7,11-HD), which is known to afford reproductive isolation between the sibling species *D. melanogaster* and *D. simulans*. A pair of alternatively spliced receptors, Or69aB and Or69aA, is tuned to *Z*4-11Al and to food olfactory cues, respectively. These receptors are co-expressed in the same olfactory sensory neurons, and feed into a neural circuit mediating species-specific, long-range communication: the close relative *D. simulans*, which shares food resources and co-occurs with *D. melanogaster*, does not respond. That Or69aA and Or69aB have adopted dual olfactory traits highlights the interplay of habitat and social signals in mate finding. These olfactory receptor genes afford a collaboration between natural and sexual selection, which has the potential to drive phylogenetic divergence.

**Significance Statement:** Volatile insect sex pheromones carry a message over a distance, they are perceived by dedicated olfactory receptors, and elicit a sequence of innate behaviours. Pheromones mediate specific mate recognition, but are embedded in and perceived together with environmental olfactory cues. We have identified the first long-range, species-specific pheromone in *Drosophila melanogaster.* A pair of spliced olfactory receptors, feeding into the same neural circuit, has developed a dual affinity to this pheromone and kairomones, encoding adult and larval food. Blends of this pheromone and kairomone specifically attract *D. melanogaster*, but not the close relative *D. simulans.* This becomes an excellent paradigm to study the interaction of social signals and habitat olfactory cues in premating reproductive isolation and phylogenetic divergence.

## Introduction

Volatile insect pheromones transmit species-specific messages and elicit long-range flight attraction. Premating communication with pheromones facilitates and accelerates mate-finding, and reduces predation risk and energy expenditure, which is particularly adaptive in short-lived insects (1,2).

Sexual communication subserves mate-finding and ultimately reproduction. Female insects search for larval food and oviposition sites soon after mating, and both sexes forage to offset the nutritional cost of reproduction. The search for mates and food is accordingly interconnected, and so is the response to sex and habitat olfactory signals. The sensory drive hypothesis predicts that mate recognition in animals evolves during niche adaptation and that premating sexual communication involves olfactory specialization to both social signals and habitat cues (3,4).

Pheromones are released into an atmosphere that is filled with environmental, habitat-related olfactory cues, some of which signal mating sites and food sources. The response to sex pheromones and food or habitat odourants (kairomones) is under sexual and natural selection, respectively. Pheromones and kairomones are always perceived as an ensemble in a natural context and this leads to an interaction of sexual and natural selection during adaptive divergence of sexual signaling, which is thought to facilitate premating reproductive isolation (3–7).

Olfactory sexual communication is studied at cellular and molecular resolution in the fruit fly *Drosophila melanogaster*, but volatile pheromones encoding species-specific, long-range mate recognition have not yet been found. *Drosophila* is attracted to yeast and fruit odorants for feeding, mating and oviposition (8–10) and the interconnection between perception of pheromones and food semiochemicals is a current research theme (11, 12). For example, the male-produced sex pheromone cVA and food stimuli are integrated to coordinate feeding, courtship behavior and oviposition site selection (13–16). Perception of cVA is a current and outstanding paradigm for studying the molecular and neuronal logic of innate, olfactory-mediated reproductive behavior (15, 17, 18). cVA and other known olfactory pheromones are active during courtship, and since they are all shared with other *Drosophila* species, they cannot account for species-specific communication (19, 20, 21).

*D. melanogaster* and *D. simulans* are sibling species with no gene flow between them (22). Interspecific matings of *D. melanogaster* with *D. simulans*, or other closely related species are inhibited by the female-produced cuticular hydrocarbon (*Z*,*Z*)-7,11-heptacosadiene (7,11-HD), which is perceived through gustatory receptors at close range (23–25). 7,11-HD or other non-volatile cuticular hydrocarbons convey species-specificity, but they are not volatile and cannot account for long-range communication and flight attraction. This raises the question whether *Drosophila* uses, in addition, volatile pheromone signals that mediate specific mate recognition at a distance.

We have identified the first long-range, species-specific pheromone in *D. melanogaster.* A pair of spliced olfactory receptors, feeding into the same neural circuit, has developed a dual affinity to this pheromone and to environmental semiochemicals, encoding adult and larval food. A blend of this pheromone and a food odourant specifically attracts *D. melanogaster*, but not the close relative *D. simulans.* This becomes an excellent paradigm for studying the interaction of social signals and habitat olfactory cues in premating reproductive isolation and phylogenetic divergence.

## Results

### *Drosophila melanogaster* females produce a suite of volatile aldehydes

A focus in *Drosophila* pheromone research has been on cuticular hydrocarbons, which are active during close-range courtship. Our scope was to investigate volatile compounds encoding long-range communication. We therefore collected volatile compounds released by *D. melanogaster* flies in a glass aeration apparatus and found 16 aliphatic aldehydes, according to chemical analysis by gas chromatography-mass spectrometry (GC-MS). Males and females shared saturated aldehydes with a carbon chain length of C7 to C18, but mono-unsaturated aldehydes were released by females only (Fig. 1*A*; Table 1). The most abundant compound was identified as (*Z*)-4-undecenal (*Z*4-11Al) and synthesized.

**Figure 1.**
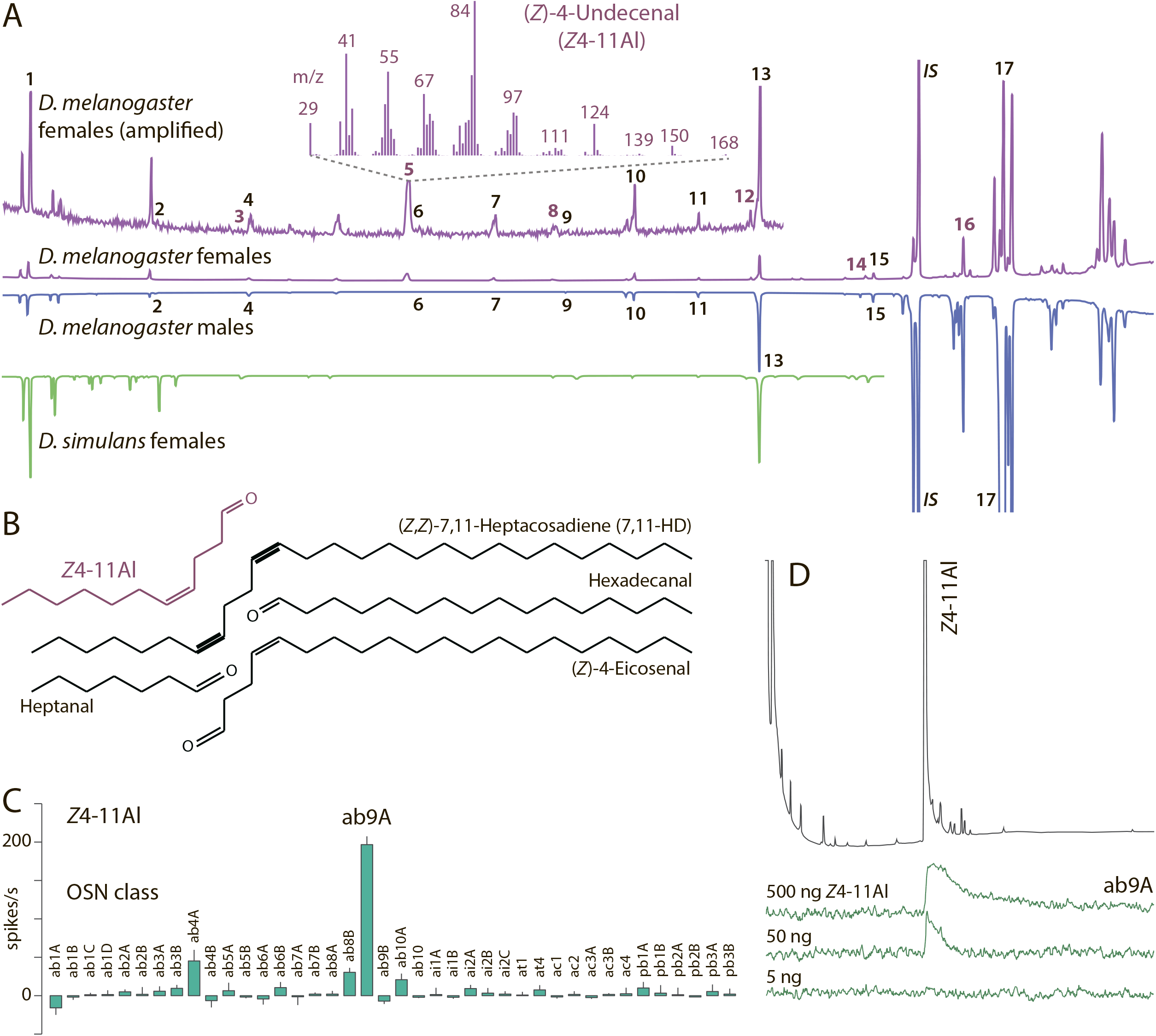
Headspace analysis of *Drosophila* females and males by GC-MS and electrophysiological screening of the candidate pheromone compound Z4-11A! on male antennae. **(a)** Chromatograms of headspace collections from *D. melanogaster* females (lilac traces; upper trace: amplified signal; lower trace: entire chromatogram), males (blue trace), and *D. simulans* females (green trace). The headspace of *D. melanogaster* females contains 16 yet undescribed compounds: heptanal (*1*), octanal (*2*), (*Z*)4-nonenal (*3*), nonanal (*4*), (*Z*)4-undecenal (*Z*4-11Al) (*5*), undecanal (*6*), dodecanal (*7*), (*Z*)4-tridecenal (*8*), tridecanal (*9*), tetradecanal (*10*), pentadecanal (*11*), (*Z*)4-hexadecenal (*12*), hexadecanal (*13*), (*Z*)4-octadecenal (*14*), octadecanal (*15*) and (*Z*)4-eicosenal (*16*) (see Table 1). Female-specific compounds are coloured, the most abundant cuticular hydrocarbon, 7-tricosene (*17*) is shown for reference, the internal standard (IS) was heptadecyl acetate. Inset: mass spectrum of the most abundant female-specific compound *Z*4-11Al. **(b)** Oxidation of the most abundant female cuticular hydrocarbon (*Z*,*Z*)-7,11-heptacosadiene (7,11-HD), affording two saturated and two unsaturated aldehydes, heptanal, hexadecanal, *Z*4-11Al and (*Z*)4-eicosenal. **(c)** Single sensillum recordings (SSR) from all *D. melanogaster* olfactory sensory neurons (OSNs) with *Z*4-11Al (error bars show SEM; *n* = 5). **(d)** SSR coupled to GC (GC-SSR), showing a response of ab9A to three different amounts of *Z*4-11Al.

**Table 1.** Saturated and unsaturated aldehydes found in headspace collections of *D. melanogaster* (Dalby) females and males.

| Compound | Females<br>(%±SD, n=10) | Males<br>(%±SD, n=10) |
| --- | --- | --- |
| heptanal | 3.1 ± 0.8 | 9.9 ± 4.8 |
| octanal | tr **a** | 0.2 ± 0.5 |
| nonanal | 4.1 ± 0.9 | 8.7 ± 7.9 |
| (Z)-4-nonenal | tr | - **b** |
| undecanal | tr | - |
| (Z)-4-undecenal | 23.3 ± 1.8 | - |
| dodecanal | 7.9 ± 1.3 | - |
| tridecanal | tr | - |
| (Z)-4-tridecenal | 0.4 ± 0.9 | - |
| tetradecanal | 11.2 ± 0.8 | 4.9 ± 4.1 |
| pentadecanal | 3.2 ± 0.9 | 2.4 ± 2.1 |
| hexadecanal | 27.8 ± 3.3 | 69.1 ± 9.7 |
| (Z)-4-hexadecenal | 2.9 ± 0.3 | - |
| octadecanal | 4.7 ± 1.0 | 4.4 ± 3.9 |
| (Z)-4-octadecenal | 3.0 ± 0.6 | - |
| (Z)-4-eicosenal | 5.6 ± 1.7 | - |

GC-MS analysis showed that *Z*4-11Al was present also in cuticular extracts of females, although in lower amounts (0.27 ± 0.12 ng/female, *n* = 5) than in headspace collections (3.0 ± 0.81 ng/female, *n* = 10; *P*<0.01 Mann-Whitney test). Cuticular profiles of *Drosophila* flies have been investigated, but *Z*4-11Al or other aldehydes have not been reported (19, 26, 27).

The sister species *D. simulans* did not release *Z*4-11Al, nor other monounsaturated aldehydes (Fig. 1*A*). Unlike *D. melanogaster, D. simulans* does not produce (*Z*,*Z*)-7,11-heptacosadiene (7,11-HD) (23). This led us to hypothesize that the production of monounsaturated aldehydes with a double bond in position 4 was linked to oxidation of diunsaturated cuticular hydrocarbons. Oxidation of 7,11-HD is expected to generate two saturated aldehydes, heptanal and hexadecanal, and two unsaturated aldehydes, *Z*4-11Al and (*Z*)-4-eicosenal (Fig. 1*B*). This was experimentally verified by applying 100 ng synthetic 7,11-HD to a glass vial. After 60 min, 1.92 ± 0.42 ng *Z*4-11Al were retrieved (*n* = 3). Based on the cuticular hydrocarbon profile of *D. melanogaster* (24), 26 aldehydes are expected to be formed by oxidation, 16 of which were found in our headspace analysis (Table 1), others may have been below detection level.

Next, single sensillum electrophysiological recordings (SSR) from all basiconic, trichoid, coeloconic, and intermediate olfactory sensilla in *D. melanogaster* (Fig. 1*C*) and GC-coupled SSR recordings (GC-SSR) from ab9 sensilla (Fig. 1*D*) showed that *Z*4-11Al strongly activates ab9A olfactory sensory neurons (OSNs). A weaker response from ab4A neurons to *Z*4-11Al probably reflects the sensitivity of Or7a (expressed in ab4A) to aldehydes, such as the leaf volatile (*E*)-2-hexenal or bombykal, a lepidopteran pheromone compound (28–30).

### The olfactory receptor Or69aB responds to *Z*4-11Al

ab9A OSNs express the olfactory receptor (Or) Or69a (31). We therefore screened ab9A OSNs with known ligands of Or69a (30) and *Z*4-11Al. In the *D. melanogaster* strains Canton-S and Zimbabwe, the monoterpene (*R*)-carvone elicited the strongest response from ab9A, although the response to *Z*4-11Al was not significantly different. In *D. simulans, Z*4-11Al elicited a significantly lower response than (*R*)-carvone (Fig. 2*A).*

**Figure 2.**
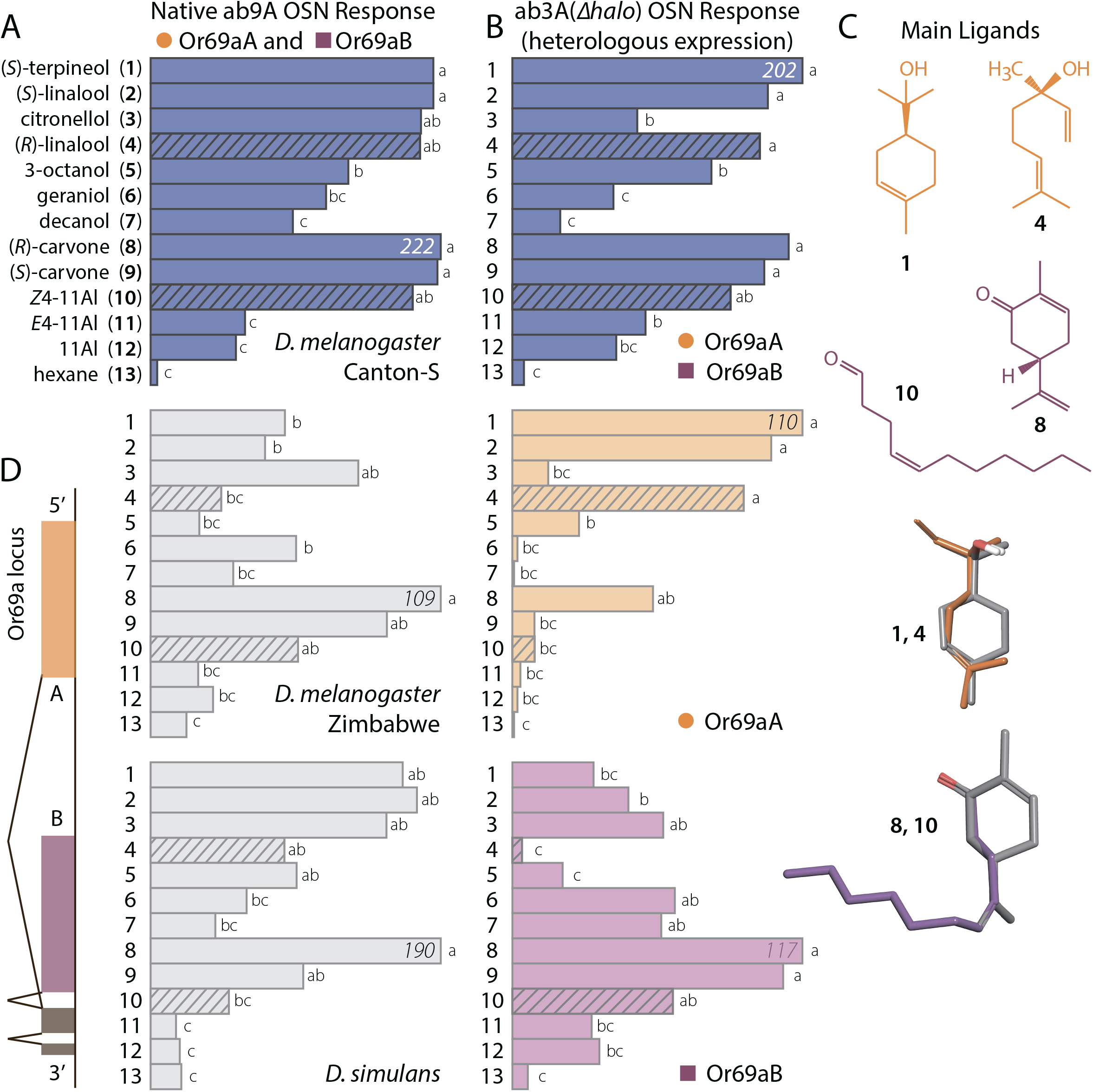
SSR-Response of Or69a splice variants to ten odorants, in native ab9A OSNs and in ab3A OSNs, following heterologous expression. **(a)** SSR from ab9A OSNs, in *D. melanogaster* (Canton-S, Zimbabwe) and *D. simulans* males, which natively express both splice variants Or69aA and Or69aB. Responses were normalized to the most active odorant. **(b)** SSR from ab3A OSNs in *D. melanogaster*, heterologously expressing Or69aA and Or69aB, together and singly. Test panel includes the known most active ligands for Or69a (27) and three aldehydes. Cross-hatched bars indicate behaviorally active compounds (Fig. 3). Bars followed by different letters indicate statistically significant differences for each fly type (*P*<0.05; Mann-Whitney test, *n* = 5 for ab9A, *n* = 10 for ab3A). **(c)** Key ligands for Or69aA, (*S*)-terpineol (1) and (*R*)-linalool (*4*), and for or Or69aB, (*R*)-carvone (*8*) and *Z*4-11Al (*10*). Alignment of these ligands illustrates shared structural motifs. **(d)** Alternative splicing of Or69a, where coloured boxes A and B show unique exons encoding the splice products; dark boxes show shared exons, generating co-expression of Or69aA and Or69aB in the same neurons in ab9 sensilla in *D. melanogaster.*

The Or69a gene encodes two proteins, Or69aA and Or69aB, as a result of alternative splicing (Fig. 2*D*), which occurred prior to the split of the *D. obscura* and *D. melanogaster* groups (32, 33). Heterologous co-expression of both Or69a splice variants in ab3A (*Δhalo*) empty neurons (34) produced a response similar to native ab9A OSNs; whereas individual expression revealed distinct response profiles for Or69aA and Or69aB (Fig. 2 *A* and *B*). Or69aB responds best to both isomers of carvone, followed by *Z*4-11Al.

Carvone and *Z*4-11Al, which seem unalike at first glance, share a structural motif, a carbonyl functional group with an equidistant double bond in position 4 (Fig. 2 *B* and *C*). Different ligands, upon binding to the same Or, are thought to adopt a complementary bioactive conformation. The strain energy required for any compound to assume a steric conformation that aligns with an active ligand should typically not exceed 5 kcal/mol (35). Conformational analysis showed that *Z*4-11Al aligns with (*R*)-carvone, which elicited the strongest Or69aB response, at a strain energy cost of only 1.5 kcal/mol. Or69aA, on the other hand, is tuned to terpenoid alcohols and responded significantly less to *Z*4-11Al. The most active ligands (*S*)-terpineol, (*S*)-and (*R*)-linalool, which again share the functional group and a double bond in position 4, align at 3.0 kcal/mol (Fig. 2 *B* and C). Conformational analysis confirms that the most active ligands for Or69aA and Or69aB, respectively, are structurally related.

### Z4-11Al elicits upwind flight attraction in *D. melanogaster*, but not in *D. simulans.*

*Z*4-11Al elicited upwind flight and landing at the source, in cosmopolitan Dalby and Canton-S strain *D. melanogaster* males and females. In contrast, males of the Zimbabwe strain and *D. simulans* were not attracted (Fig. 3 *A* and *B*). This shows that *Z*4-11Al, in addition to its precursor 7,11-HD (Fig. 1) participates in sexual isolation between *D. melanogaster* and its sister species *D. simulans* (23), and between cosmopolitan and African *D*. *melanogaster* strains (36–38).

**Figure 3.**
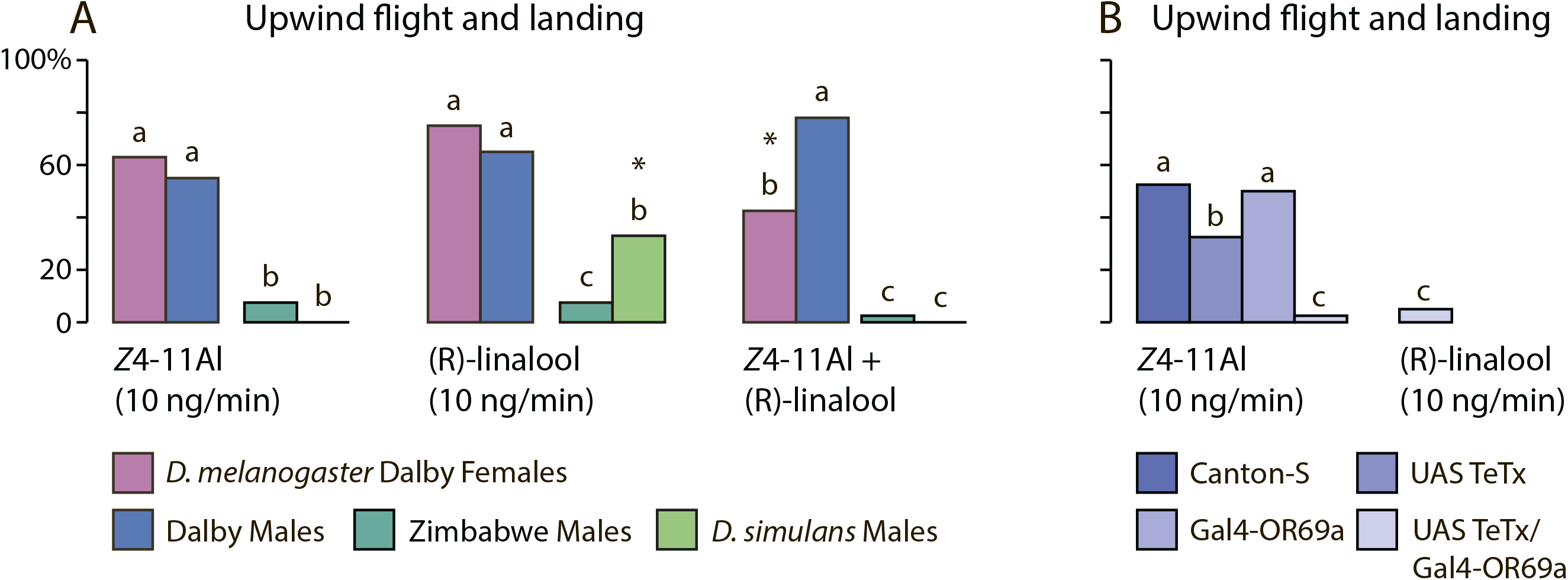
*Z*4-11Al mediates long range attraction in *D. melanogaster.* **(a)** Upwind flights to 10 ng/min of Z4-11A! and (*R*)-linalool, followed by landing at the source, in *D. melanogaster* (Dalby) males and females, in *D. melanogaster* (Zimbabwe) males and *D. simulans* males. Lower case letters indicate statistical differences between test insect strains and species, for each treatment. Asterisks indicate significant differences between treatments (*n* = 40, *P*<0.001; binomial GLMs followed by post-hoc Wald pairwise comparison tests).**(b)** Upwind flights to 10 ng/min of *Z*4-11Al and (*R*)-linalool in *D. melanogaster* (CantonS) males expressing a tetanus toxin in OSNs expressing Or69a, and in the parental lines. Letters indicate statistical differences within treatments (*n* = 40, *P*<0.001; binomial GLM, followed by *Post-hoc* Wald pairwise comparison tests).

Moreover, admixture of *Z*4-11Al eliminated *D. simulans* attraction to the yeast volatile (*R*)-linalool (39) (Fig. 3*A*). *D. melanogaster* and its sister species *D. simulans* co-occur in the same habitat and use partially overlapping food resources. The very rarely occurring hybrid matings are sterile; the antagonistic interaction between pheromone and food stimulus is therefore adaptive and substantiates a contributing role of Or69a in reproductive isolation. A tentative explanation for the significantly reduced attraction response of *D. melanogaster* females to the blend of *Z*4-11Al and linalool, compared to males (Fig. 3*A*) is that females would avoid food resources and rendezvous sites overcrowded with other females.

The response magnitude of wild-type flies to *Z*4-11Al released at a rate of 10 ng/min was similar to the upwind flight response to vinegar headspace, when the main compound acetic acid was released at a 170-fold amount (Fig. 3*A*; 36). This illustrates the high responsiveness and sensitivity of *D. melanogaster* males and females to *Z*4-11Al.

Finally, we used tetanus toxin mutants to verify that Or69a encodes *Z*4-11Al. Upwind flight attraction to *Z*4-11Al or linalool was significantly reduced when Or69a OSNs were disrupted (Fig. 3*B*). In summary, *Z*4-11Al is a powerful attractant that enables specific mate recognition at a distance. Its heterospecific role was conserved even in blends with the food attractant linalool.

## Conclusions

*Z*4-11Al is the first species-specific, long-range sex pheromone of *D. melanogaster.* It is produced by females and perceived by Or69aB in both sexes. The precursor of *Z*4-11Al is the cuticular hydrocarbon 7,11-HD, which mediates isolation between *D. melanogaster* and its sister species *D*. *simulans* during courtship (23–25).

In addition to pheromone, Or69aB and its twin receptor Or69aA bind kairomonal terpenoids, such as linalool or terpineol, which are found in both fruit and yeast headspace. Citrus peel, a preferred oviposition substrate (8), and baker’s yeast which grows on ripe fruit and elicits fly attraction and oviposition (9) are sources of all main ligands of Or69aA and Or69aB (39, 41).

Combined pheromone and food odour tuning in the two Or69a splice variants underscores the tie between sexual and natural selection during the evolution of specific mate communication (3, 4). Flies releasing pheromone tint pervading habitat and food odorants and thus shape and foreground a communication channel that facilitates mate finding. This is particularly adaptive in *Drosophila* when mating sites, fruit and berries, are abundant and widely spread.

A twin receptor, emerging from the Or69a splice event (32,33), facilitated adaptive changes in ligand tuning, without compromising the established functional role of the Or69a channel. Functional divergence has apparently been biased towards structurally related ligands (Fig. 2 *B* and *C*; 30) and ecologically relevant odorant signals. This is expected; conservative diversification of the splice variants is constrained to a behavioral theme, since both Ors feed into one OSN.

Olfactory representations of other *Drosophila* Ors involved in food and pheromone perception project through separate channels to the LH, where third-order neurons partially overlap and integrate (15). In stark contrast, Or69a is the first olfactory gene known to encode dual olfactory traits. Or69aA and Or69aB co-express in the same ab9A OSNs (31) and thus achieve a coordination of mating and food stimuli already in first order neurons, at the antennal periphery. This makes Or69a a target for selection during phylogenetic divergence. The tuning range of Ors evolves more rapidly than hardwired neural circuits in higher brain centres (42) and selection pressure is further relaxed following a splice event. Differential tuning of Or69a in closely related cosmopolitan and African strains of *D. melanogaster* (Fig. 2 *A*) corroborates this idea.

Tuning changes in the two splice forms of Or69a are restricted with respect to the behavioral and ecological role of their ligands, since they both feed into a neural circuit mediating sexual and habitat attraction. The two splice forms provide, on the other hand, degrees of freedom during adaptive divergence, since they allow fly populations to adopt new kairomone or pheromone signals; alteration of either one produces a new communication channel. Reproductive isolation may arise as a byproduct (43–45) and the Or69a gene therefore has the potential to drive speciation (46, 47). Species in the *D. melanogaster* and *D. obscura* groups provide a rich substrate for studying Or ligand evolution and its consequences on disruptive selection on ecological interactions and mate choice.

## Methods

### Insects

Canton-S, Zimbabwe (S-29; Bloomington #60741) and Dalby-HL (Dalby, Sweden) (48) strains of *D. melanogaster* were used as wild type flies for behavioral experiments. Canton-S was used for comparison with knockouts of the same background. Further tests were done with the sister species *D. simulans.*

We used the Or69a-Gal4/UAS TeTx, tetanus toxin knockout line to verify the role of Or69a in flight attraction to *Z*4-11Al. Canton-S/UAS TeTx (Bloomington #28838 and 28997) and Canton-S/Or69a-Gal4 (Bloomington #10000) were used as parental controls.

Flies were reared on a standard sugar-yeast-cornmeal diet at room temperature (19 to 22°C) under a 16:8-h L:D photoperiod. Newly emerged flies were anesthetized under CO_2_ and sexed under a dissecting microscope. Virgin flies were identified by the presence of meconium, and were kept together with flies of the same sex. Flies were kept in 30-ml Plexiglas vials with fresh food. Experiments were done with 3-to 5-d-old flies.

### Chemicals

(*Z*)-4-undecenal (*Z*4-11Al) and (*E*)-4-undecenal (*E*4-11Al) were synthesized. A short description follows below, for a complete account of the chemical synthesis, see Supplemental Information.

(*Z*)-4-Undecenoic acid was synthesized via a modified version of Wube et al. (49) in 80% stereoisomeric purity. Esterification under acidic conditions with sulfuric acid in methanol resulted in 80% *Z*-isomer and a 93% yield over two steps. Stereoisomeric purity was controlled with NMR and GC-FID by comparing analysis for acid and ester, the appearance of a small quartet, in the NMR spectra, at 1.96 indicates the presence of *E*-isomer. Gas chromatographic separation on a polar Varian factorFOUR vf-23ms of *Z*-and *E*-ester proved that the stereochemistry was not affected by the acidic conditions during esterification. Methyl (*Z*)-4-undecenoate was purified on regular silica gel and on silver nitrate impregnated silica gel to obtain a stereoisomeric purity of 98.6%. Methyl (*Z*)-4-undecenoate was reduced to (*Z*)-4-undecenol with lithium aluminum hydride in diethylether and oxidized to *Z*4-11Al with Dess-Martin periodinane in dichloromethane.

A modified version of Virolleaud’s (50) metathesis was used to produce (*E*)-4-undecenoic acid in a 56% yield (87.5% of the *E*-isomer). (*E*)-4-undecenoic acid was esterified under the same conditions as the (*Z*)-acid, without isomerisation of the double bond (according to GC-FID and ^1^H-NMR). The methyl-(*E*)-4-undecenoate was reduced to the alcohol with lithium aluminum hydride in diethylether and purified on silver nitrate impregnated silica gel to obtain a purity of 99.8% of the (*E*)-isomer, which was oxidized with Dess-Martin periodinane in dichloromethane to obtain *E*4-11Al.

Commercially available compounds were: (*R*)-carvone (97% chemical purity, CAS #648540-1, Firmenich), (*S*)-carvone (98%, CAS #2244-16-8, Firmenich), (*S*)-terpineol (97%, CAS #10482-56-1, Aldrich), (*S*)-linalool (97%, CAS #126-91-0, Firmenich), (*R*)-linalool (97%, CAS #126-90-9, Firmenich), citronellol (99%, CAS #106-22-9, Aldrich), geraniol (98%, CAS #106-24-1, Aldrich), 3-octanol (99%, CAS #589-98-0, Aldrich), decanol (99%, CAS #112-30-1, Fluka), 11-Al (99%, CAS #112-44-7, Aldrich).

### Odor collection and chemical analysis

Twenty *D. melanogaster* (Dalby), *D. melanogaster* (Canton) (*n* = 10) or 20 *D. simulans* (*n* = 10) unmated female or unmated male flies were placed in a glass aeration apparatus designed for collection of airborne pheromone (effluvia) (51). The flies were held in a glass bulb with a narrow open outlet (ø 1 mm), which prevented them from escaping. A charcoal-filtered air flow (100 mL/min) passed over the flies during 75 min. Fly effluvia were collected on the glass surface, breakthrough was monitored by attaching a 10-cm glass capillary (ø 1 mm) attached to the outlet. After 75 min, flies were removed, 100 ng of heptadecyl acetate (internal standard) was deposited in the glass bulb, which was then rinsed with 50 μl hexane, the solvent was concentrated to 10 μl in Francke vials.

Cuticular extracts (*n* = 5) were obtained by dropping 20 *D. melanogaster* females for 5 min in 400 μl hexane containing 100 ng heptadecyl acetate. After 5 min, the extracts were transferred to Francke vials and concentrated to 10 μl before analysis. Fly extracts and volatile collections were stored at −20°C.

Oxidation of (*Z*,*Z*)-7,11-heptacosadiene (7,11-HD) was analysed by dropping 100 ng synthetic 7,11-HD into a 1.5-mL glass vial, at 19°C. Vials were rinsed with 10 μl of hexane, which contained 100 ng heptadecyl acetate as an internal standard, after 15, 30, 45, 60 and 75 min (*n* = 3).

Samples were analysed by combined gas chromatography and mass spectrometry (GC-MS; 6890 GC and 5975 MS, Agilent technologies Inc., Santa Clara, CA, USA). Two μl were injected (injector temperature 225°C) splitless (30 s) into fused silica capillary columns (60 m × 0.25 mm), coated with HP-5MS UI (Agilent Technologies Inc., d_f_ = 0.25 μm) or DB-wax (J&W Scientific, Folsom, CA, USA, d_f_ = 0.25 μm), that were temperature-programmed from 30 to 225°C at 8°C/min. Helium was used as mobile phase at 35 cm/s. The MS operated in scanning mode over m/z 29–400. Compounds were tentatively identified based on their mass spectra and Kovats retention indices, using custom and NIST (Agilent) libraries, followed by comparison with authentic standards.

### Behavioural assays

Upwind flight behavior was observed in a glass wind tunnel (30 × 30 × 100 cm) equipped with a piezo sprayer (40). The flight tunnel was lit diffusely from above, at 13 lux, temperature ranged from 22 to 24°C, relative humidity from 38% to 48% and charcoal filtered air, at a velocity of 0.25 m/s, was produced by a fan (Fischbach GmbH, Neunkirchen, Germany). Compounds were delivered from the centre of the upwind end of the wind tunnel via a piezo-electric micro-sprayer (52). Forty flies were flown individually to each treatment. “Attraction” was defined as upwind flight, directly from a release tube at the end of the tunnel over 80 cm towards the odor source, followed by landing. Unmated fed, 3-d-old Dalby wild-type males and females, *D. melanogaster* Zimbabwe strain males and *D. simulans* males were flown towards (*Z*)-4-undecenal (released at 10 ng/min), (*R*)-linalool (10 ng/min) and the blend of (*Z*)-4-undecenal and (*R*)-linalool (10 ng/min, each).

### Heterologous expression of Or69aA and Or69aB

Or69aA and Or69aB receptors were cloned from antennae of *D. melanogaster*, Dalby line (53). Briefly, cDNA was generated from RNA extracts of antennae of 100 males and females using standard procedures. Or69a variants were PCR amplified with the following primers: Or69aA_5′: GTCATAGTTGAAACCAGGATGCAGTTGC, Or69aB_5′: ATAATTCAGGACTAGATGCAGTTGGAGG, Or69aAB_3′: TGCACTTTTGCCCTTTTATTTAAGGGAC.

The splice variants were amplified with unique 5′ primers and a common 3′ primer, reflective of genomic structure at this locus. These primers encompass the entire open reading frame of the receptor variants, and are located partially upstream and downstream of the start and stop codons. PCR amplicons were gel-purified and cloned into the pCR8/GW/Topo-TA Gateway entry vector (Thermo-Fisher Scientific, Waltham, MA, USA) according to standard procedure, with vector inserts sequenced to confirm fidelity of Or sequence. Or inserts were subsequently transferred to pUAS.g-HA.attB (54) with LR Clonase II enzyme (Thermo-Fisher Scientific), according to manufacturers protocol; vector inserts were sequenced to confirm fidelity of Or sequence.

Mini-prep purified pUAS.g-HA.attB plasmid with Or69aA or Or69aB insert were delivered to Best Gene Inc. (Chino Hills, CA USA) for generation of transgenic *D. melanogaster* flies. Using the PhiC31 targeted genomic-integration system (54) vectors with Or69aA or Or69aB were injected into the following fly strain, for integration on the 3^rd^ chromosome: M{3xP3-RFP.attP}ZH-86Fb (with M{vas-int.Dm}ZH-2A) (Bloomington *Drosophila* Stock Number: 24749). For expression of single receptor variants in the empty neuron system, Or69a transgenes were crossed into the *Δhalo* background to give genotype: w; *Δhalo*/Cyo; UAS-DmelOr69a(A or B), and these flies were crossed to flies with genotype: w; *Δhalo*/Cyo; DmelOr22a-Gal4, as described previously (53). Experimental electrophysiology assays were performed on flies with genotype: w; *Δhalo* UAS-DmelOr69a(A or B)/DmelOr22a-Gal4.

For co-expression of Or69aA and Or69aB in the same empty neurons, a second fly-line with Or69aB was generated with Or69aB present on the X-chromosome. The same UASg-HA.attB:Or69aB plasmid generated previously was injected into the following fly strain: y,w, P{CaryIP}su(Hw)attP8 (Bloomington *Drosophila* Stock Number: 32233). The Or69aB transgene was crossed into the DmelOr22a-Gal4 line in *Δhalo* background to give genotype: UAS-DmelOr69aB; *Δhalo*/Cyo; DmelOr22a-Gal4; these flies were crossed to flies with genotype: w; *Δhalo*/Cyo; UAS-DmelOr69aA. Experimental electrophysiology assays were performed on flies with genotype: UAS-DmelOr69aB/w; *Δhalo*; UAS-DmelOr69aA/DmelOr22a-Gal4.

### Conformational analysis

MacroModel version 11.0 (Schrodinger LLC, New York, NY, USA) in the Maestro Version 10.4.017 were used to build, minimize and to perform conformational analysis of *Z*4-11Al, (*R*)-carvone, (*S*)-terpineol and (*R*)-linalool, using default settings (OPLS3 as force field, water as the solvent and mixed torsional/low-mode sampling method). The assumed bioactive conformations of the conformationally more flexible compounds, *Z*4-11Al and (*R*)-linalool, were based on the positon of the shared functional groups in the conformationally more restricted compounds, (*R*)-carvone and (*S*)-terpineol. The carbonyl and the double bond atoms were kept fixed during minimization of the proposed bioactive conformation of *Z*4-11Al; the alcohol functional group and the double bond were kept fixed in (*R*)-linalool. The strain energies, the energy cost for adopting proposed bioactive conformations, were then calculated as the difference between the lowest energy conformations and the assumed bioactive conformation.

### Electrophysiological recordings

Single sensillum recordings (SSR) were done as described earlier (19). Unmated males were restrained in 100-μl pipette tips, with half of the head protruding, the third antennal segment or palps were placed on a glass microscope slide and held by dental wax. For the initial screening, all basiconic, trichoid, coeloconic, and intermediate sensilla (31) were localized in *D. melanogaster* (Canton-S strain) males, under a binocular at 1000x magnification. Further recordings were made from small basiconic ab9 sensilla, in *D. melanogaster* (Canton-S and Zimbabwe strains) and in *D. simulans* males, and from large basiconic ab3 sensilla in mutant *D. melanogaster*, where Or69aA and Or69aB were heterologously expressed (see above).

Tungsten electrodes (diameter 0.12 mm, Harvard Apparatus Ltd, Edenbridge, United Kingdom) were electrolytically sharpened with a saturated KNO_3_ solution. The recording electrode was introduced with a DC-3K micromanipulator equipped with a PM-10 piezo translator (Märzhäuser Wetzler GmbH, Germany) at the base of the sensilla. The reference electrode was inserted into the eye. The signal from olfactory sensory neurons (OSNs) was amplified with a probe (INR-02; Syntech), digitally converted by an IDAC-4-USB (Syntech) interface, and analyzed with Autospike software v. 3.4 (Syntech). Neuron activities were recorded during 10 s, starting 2 s before odor stimulation. Neuron responses were calculated from changes in spike frequency, during 500 ms before and after odor stimulation.

Odorants were diluted in redistilled hexane, 10 μg of test compounds in 10 μl hexane were applied to filter paper (1 cm^2^), kept in Pasteur pipettes. The test panel contained the most active ligands known for Or69a (30) and several aldehydes. Diagnostic compounds for confirmation of sensillum identity were 2-phenyl ethanol (ab9) and 2-heptanone (ab3). Control pipettes contained solvent only. Puffs (2.5 ml, duration 0.5 s) from these pipettes, produced by a stimulus controller (Syntech GmbH, Kirchzarten, Germany), were injected into a charcoal-filtered and humidified airstream (0.65 m/s), which was delivered through a glass tube to the antenna.

For GC-SSR recordings, GC columns and the temperature programmes were the same as for the GC-MS analysis. At the GC effluent, 4 psi of nitrogen was added and split 1:1 in a 3D/2 low dead volume fourway-cross (Gerstel, Mühlheim, Germany) between the flame ionization detector and the antenna. Towards the antenna, the GC effluent capillary passed through a Gerstel ODP-2 transfer line, that tracked GC oven temperature, into a glass tube (30 cm × 8 mm ID), where it was mixed with charcoal-filtered, humidified air (20°C, 50 cm/s).

### Statistical analysis

Generalized linear models (GLM) with a Bernoulli binomial distribution were used to analyse wind tunnel data. Landing at source and sex were used as the target effects. *Post-hoc* Wald pairwise comparison tests were used to identify differences between treatments. For all the electrophysiological tests, differences in spike activity derived from SSRs were analyzed with Kruskall Wallis H test followed by pairwise comparisons with Mann Whitney U *post hoc* test. All statistical analysis were carried out using R (R Core Team 2013) and SPSS Version 22 (IBM Corp.).

## Acknowledgements

This study was supported by the Linnaeus initiative “Insect Chemical Ecology, Ethology and Evolution” (The Swedish Research Council Formas, SLU), Carl Tryggers Stiftelse för Vetenskaplig Forskning (Stockholm) and the Swedish University of Agricultural Sciences (LTV Faculty). EW and EH were supported by the European Regional Development Fund and the County Board of Västernorrland.

## Author Contributions

S.L., P.G.B. and P.W. conceived the study. F.B.-E. and M.S. carried out behavioural studies, F.G. and W.B.W produced empty neuron flies, F.G. and H.D. did single sensillum recordings, supervised by B.S.H, E.W. and E.H. synthesized the compound, A.-L.G. calculated the confirmational analysis, M.B., G.B. and S.L. performed chemical analysis, P.W. wrote the paper with input from all co-authors.

## Supplementary Information: Chemical Synthesis

Dry THF and dry Et_2_O was obtained from a solvent purification system (Activated alumina columns, Pure Solv PS-MD-5, Innovative technology, Newburyport, USA) and used in the reactions when dry conditions were needed. All other chemicals were used without purification. Reactions were performed under Argon atmosphere unless otherwise stated. Flash chromatography was performed on straight-phase silica gel (Merck 60, 230–400 mesh, 0.040–0.063 mm, 10–50 g/g of product mixture) employing a gradient technique with an increasing concentration (0-100 %) of distilled ethyl acetate in distilled cyclohexane. In cases of very polar products chromatography was continued with ethanol in ethyl acetate (0–20 %). Thin-layer chromatography (TLC) was performed to monitor the progress of the reaction on silica gel plates (Merck 60, precoated aluminium foil), using ethyl acetate (40 %) in cyclohexane as an eluent, and plates were developed by means of spraying with vanillin in sulfuric acid and heating at 120°C. Purity of the product was checked with gas chromatography (GC) analysis on a Varian 3300 GC instrument equipped with a flame ionization detector (FID) using a capillary column Equity-5 (30 m × 0.25 mm i.d, d_f_ = 0.25 μm, with nitrogen (15 psi) as carrier gas and a split ratio of 1:20). The oven temperature was programmed at 50°C for 5 min followed by a gradual increase of 10°C min^-1^ to reach a final temperature of 300°C. An Agilent 7890 GC equipped with a polar capillary column FactorFOUR vf-23ms (30 m × 0.25 mm i.d., d_f_ = 0.25 μm) coupled to an Agilent 240 ion-trap MS detector for separation of some isomeric intermediates. The injector was operated in split mode (1:20) at 275°C, and a helium flow rate of 1 ml min^−1^ and a transfer line temperature of 280 °C. The analyses were performed in the external ionisation configuration. EI spectra were recorded with a mass range of m/z 50–300 at fast scan rate. Nuclear magnetic resonance (NMR) spectra were recorded on a Bruker Avance 500 (500 MHz ^1^H, 125.8 MHz ^13^C) spectrometer using CDCl_3_ as solvent and internal standard.

**(*Z*)-4-Undecenoic acid**. NaHMDS (6.78 mmol, 1 M in hexane) was added dropwise, during 30 min, to a suspension of (3-carboxypropyl)triphenylphosphonium bromide (1.45 g, 3.39 mmol) in THF (25 mL). The mixture was stirred for 2 h then cooled to 0°C on ice/water bath, and heptanal (0.387g, 3.39 mmol) in THF (2.5 mL) was added slowly during 15 min. The mixture was stirred for 5 h at 0°C then allowed to reach room temperature over night. The reaction was quenched with H_2_O (20 mL) and the organic solvent was evaporated. The remaining water phase was extracted with Et_2_O (3 × 20 mL), the obtained organic phases discarded and the basic aqueous phase was acidified with HCl (2M) until pH 1 and extracted with Et_2_O (3 × 20 mL). The combined organic phases were dried over MgSO_4_ (anhydr.) and the solvent evaporated off. The obtained crude product was dissolved in pentane, cooled at −18°C and filtered to remove the precipitated OPPh_3_ followed by evaporation of the solvent to result in 0.547 g of a yellow oil (87.5% yield). ^1^H-NMR: 5.52–5.30 (m, 2H), 2.35 (m, 4H), 2.04 (q, *J* = 6.5 Hz, 1.6H, *Z*-isomer), 1.96 (q, *J* = 6.5 Hz, 0.4H, *E*-isomer), 1.37-1.19 (m, 8H), 0.89 (t, *J* = 7 Hz, 3H) ppm. The NMR data is in accordance with data previously reported (46, 52). The relationship by integration between protons at 2.04 and 1.95 indicates approximately a *E*:*Z*-ratio of 80:20 which is supported by GC-MS analysis on a Varian factorFOUR vf-23ms column. The obtained crude product was used in the next step without further purification.

**Methyl (*Z*)-4-undecenoate.** (*Z*)-4-Undecenoic acid (0.547 g, 2.97 mmol) from above was dissolved in methanol (15 mL) and 7 drops of concentrated H_2_SO_4_ were added followed by heating at 70°C over night. The mixture was allowed to reach room temperature and the methanol was evaporated and the remaining crude product was dissolved in Et_2_O (15 mL). The organic phase was washed with H_2_O (3 × 10 mL) and brine (2 × 10 mL), dried over Na_2_SO_4_ (anhydr.) and solvent evaporated resulting in 0.547 g of a yellow oil (92.8% yield). GC-MS (FactorFour vf-23ms) shows a *Z*:*E*-ratio of 80:20. ^1^H-NMR(CDCl_3_): 5.4 (m, 2H), 3.67 (s, 3H), 2.3 (m, 4H), 2.03 (q, *J* = 6.5 Hz, 1.6H, *Z*-isomer), 1.96 (q, *J* = 6.5 Hz, 0.4H, *E*-isomer), 1.33-1.21 (m, 8H), 0.89 (t, *J* = 6.5 Hz, 3H) ppm (no data found in the literature). ^13^C-NMR(CDCl_3_): 134.2, 119.9, 32.3, 31.9, 29.5, 29.3, 27.43, 22.7, 14.1 ppm. ^13^C-NMR data similar to reported (53). Proton NMR shows a 80:20 Z:E-ratio between the diastereomers. Enrichment of the *Z*-isomer on AgNO_3_ (10%) impregnated silica resulted in 63 mg of 98.6:1.4 *Z*:*E*-ratio according to GC-FID analysis on the vf-5 column as the diastereoisomeric purity was not possible to measure when using ^1^H-NMR.

**(*Z*)-4-Undecenol.** Methyl (*Z*)-4-undecenoate (63 mg, 0.32 mmol) was dissolved in Et_2_O (5 mL) and LiAlH_4_ (2 spatel tips) was added followed by stirring at room temperature for 30 min. HCl (2 M, 2 mL) was added to quench the reaction and the mixture was extracted with Et_2_O (2 × 3 mL), the combined organic layer was dried over MgSO_4_ (anhydr.) and solvent was evaporated. Purification with flash chromatography on SiO_2_ resulted in 37 mg. ^1^H-NMR(CDCl_3_): 5.43–5.32 (m, 2H), 3.67 (m, 2H), 2.16–2.10 (m, 2H), 2.08–2.02 (m, 2H), 1.69-1.60 (m, 2H), 1.39-1.22 (m, 8H), 0.89 (t, *J* = 6.5 Hz, 3H) ppm. NMR-data were similar to Kim and Hong (54) and Davis and Carlsson (55). Diastereomeric purity was checked with GC-FID before next step.

**(*Z*)-4-Undecenal.** (*Z*)-4-Undecenol (37 mg, 0.22 mmol) in DCM (3 mL) was added to Dess-Martin periodinane (0.140 g, 0.33 mmol) in DCM (0.5 mL). After 50 min, NaOH (2M, 10 mL) was added to quench the reaction. The two layers were separated and the aqueous phase was extracted with Et_2_O (3 × 10 mL), the combined organic layers were washed with NaOH (2M, 10 mL), dried over MgSO_4_ (anhydr.) and solvent was evaporated resulting in 30 mg of a yellow oil (81% yield). The crude product was purified with Kugelrohr distillation at bp 65–70 °C (1.6 mbar), resulted in 17 mg. ^1^H-NMR(CDCl_3_): 9.77 (s, 1H), 5.48–5.22 (m, 2H), 2.47 (t, *J* = 7 Hz, 2H), 2.37 (q, *J* = 7 Hz, 2H), 2.04 (q, *J* = 7Hz, 2H), 1.37-1.23 (m, 8H), 0.88 (t, *J* = 7 Hz, 3H) ppm. ^13^C-NMR(CDCl_3_): 202.1, 131.8, 127.0, 43.9, 31.8, 29.5, 29.0, 27.2, 22.6, 20.1, 14.1 ppm. Both ^1^H-and ^13^C-NMR data were in accordance with published results (56, 57). Analysis on GC-MS (FactorFour vf-23ms) resulted in a 98.6:1.4 *Z*:*E*-ratio, the *E*-isomer could not be detected by ^1^H-NMR.

**(*E*)-4-Undecenoic acid.** 4-Pentenoic acid (0.5 g, 5 mmol) and 1-octene (2.8 g, 25 mmol) was dissolved in DCM (50 mL) and Grubbs II catalyst (85 mg, 0.1 mmol) was added and the reaction was refluxed. After 7 h was a second portion of Grubbs II catalyst (85 mg, 0.1 mmol) added and the reaction refluxed for 16 h. Reaction was allowed to reach room temperature and the solvent was evaporated. The obtained crude product was dissolved in Et2O (50 mL) and filtered through a short pad of silica gel. The product was purified with flash chromatography by gradient elution (0-100% EtOAc in c-hexane followed by 0-10% EtOH in EtOAC) resulting in 0.52 g oil (56% yield). ^1^H-NMR(CDCl_3_): 5.51–5.33 (m, 2H), 2.41 (q, J=7 Hz, 2H), 2.32 (q, *J* = 7 Hz, 2H), 2.04 (q, *J* = 6.5 Hz, 0.25 H, *Z*-isomer), 1.97 (q, *J* = 6.5 Hz, 1.75H, *E*-isomer), 1.37-1.22 (m, 9H), 0.88 (t, *J* = 7.5 Hz, 3H) ppm. The relation between proton at 2.04 and 1.97 reveals a 87.5:12.5 *Z*:*E*-ratio. The isolated product was used in the next step without further purification.

**Methyl (*E*)-4-Undecenoate.** (*E*)-4-Undecenoic acid (0.52 g, 2.82 mmol) was dissolved in methanol (25 mL) and a catalytic amount H_2_SO_4_ was added and the mixture was refluxed over night. After evaporation of the solvent the crude product was dissolved in Et_2_O (10 mL) and washed with H_2_O (20 mL). The aqueous phase was extracted with Et_2_O (2 × 25 mL), the combined organic layer was washed with H_2_O (20 mL) and brine (20 mL), dried over MgSO_4_ (anhydr.) and evaporation of solvent resulted in 0.439 g (78% yield). ^1^H-NMR(CDCl_3_): 5.51–5.33 (m, 2H), 3.67 (s,3H), 2.40–2.27 (m, 4H), 1.96 (q, *J* = 6.5 Hz, 2H), 1.38-1.21 (m, 8H), 0.88 (t, *J* = 6.5 Hz,3H) ppm. Purification with flash chromatography resulted in 0.401g (71.7% yield). GC-FID showed the same stereoisomeric ratio as for the acid above.

**(*E*)-4-Undecen-1-ol.** LiAlH_4_ (0.055 g, 1.46 mmol) was added to methyl (*E*)-4-undecenoate (0.145 g, 0.73 mmol) dissolved in Et_2_O (5 mL). After 30 minutes was HCl (2 M, 5 mL) added to quench the reaction. The acidic water phase was extracted with Et_2_O (3 × 10 mL) and the combined organic layers were dried over MgSO_4_ (anhydr.) and evaporation of solvent resulted in 0.104 g (99% yield). Enrichment of the *E*-isomer with medium pressure liquid chromatography (MPLC) on AgNO_3_ (10% impregnated) silica resulted in 30 mg of a clear oil (>99.8 % E). ^1^H-NMR(CDCl_3_): 5.43 (m, 2H), 3.65 (m, 2H), 2.08 (q, *J* = 7 Hz, 2H), 1.97 (q, *J* = 7 Hz, 2H), 1.63 (pent, 2H), 1.35-1.21 (m, 9H), 0.88 (t, *J* = 6.5 Hz, 3H) ppm. ^13^C-NMR(CDCl_3_): 134.4, 131.3, 129.4, 62.6, 32.6, 32.5, 31.7, 29.6, 29.5, 28.9, 28.8, 22.6, 14.1 ppm. All NMR-data were in accordance with previous published data (58).

**(*E*)-4-Undecenal**. Dess-Martin Periodinane (0.110 g, 0.26 mmol) was added to (*E*)-4-undecen-1-ol (0.030 g, 0.22 mmol) in DCM (4 mL). NaOH (2 M, 10 mL) was added after 1 h to quench reaction. The aqueous phase was extracted with Et_2_O (3 × 10 mL) and the combined organic layers were dried over MgSO_4_ (anhydr.) and evaporation of the solvent resulted in 30 mg (98% yield). Purification of the crude with Kugelrohr distillation at 65°C (2 mbar) resulted in 10 mg of product (33% yield, 97% chemical purity, 3% undecenal). ^1^H-NMR(CDCl_3_): 9.76 (t, *J* = 1.5 Hz, 1H), 5.50–5.36 (m, 2H), 2.48 (d of t, *J* = 7.5, 1.5 Hz, 2H), 2.33 (q, *J* = 7 Hz, 2H), 1.97 (q, *J* = 6.5 Hz, 2H), 1.32-1.19 (m, 8H), 0.87 (t, *J* = 6.5 Hz, 3H). ^13^C-NMR (CDCl_3_): 202.5, 132.2, 127.6, 43.6, 32.5, 31.7, 29.4, 28.8, 25.2, 22.6, 14.1. The NMR-data were in accordance with published data (57, 58).

(*Z*)-4-Undecenoic acid was synthesized via a modified version of Wube and Hüfner (46) in 80% stereoisomeric purity. Esterification under acidic conditions with sulfuric acid in methanol resulted in 80% *Z*-isomer and a 93% yield over two steps. Stereoisomeric purity was controlled with NMR and GC-FID by comparing analysis for acid and ester, the appearance of a small quartet, in the NMR spectra, at 1.96 indicates the presence of *E*-isomer. Gas chromatographic separation on a polar Varian factorFOUR vf-23ms of *Z*-and *E*-ester proved that the stereochemistry was not affected by the acidic conditions during esterification. Methyl-(*Z*)-4-undecenoate was purified on regular silica gel and on silver nitrate impregnated silica gel to obtain a stereoisomeric purity of 98.6 %. Methyl-(*Z*)-4-undecenoate was reduced to (*Z*)-4-undecenol with lithium aluminum hydride in diethylether and oxidized to (*Z*)-4-undecenal with Dess-Martin periodinane in dichloromethane.

A modified version of Virolleaud’s (47) metathesis was used to produce the (*E*)-4-undecenoic acid and in a 56 % yield (87.5% of the *E*-isomer). (*E*)-4-undecenoic was esterified under the same conditions as the (*Z*)-acid and once again there was no isomerisation of the double bond (according to GC-FID and ^1^H-NMR. The methyl-(*E*)-4-undecenoate was reduced to alcohol with LiAlH_4_ in diethylether and purified on silver nitrate impregnated silica gel to obtain a purity of 99.8 % of the (*E*)-isomer, which was oxidized with Dess-Martin periodinane in dichloromethane to obtain the wanted (*E*)-4-undecenal.

